# Intergenerational epigenetic inheritance in reef-building corals

**DOI:** 10.1101/269076

**Authors:** Yi Jin Liew, Emily J. Howells, Xin Wang, Craig T. Michell, John A. Burt, Youssef Idaghdour, Manuel Aranda

## Abstract

The notion that intergenerational or transgenerational inheritance operates solely through genetic means is slowly being eroded: epigenetic mechanisms have been shown to induce heritable changes in gene activity in plants ^1,2^ and metazoans ^1,3^. Inheritance of DNA methylation provides a potential pathway for environmentally induced phenotypes to contribute to evolution of species and populations ^1–4^. However, in basal metazoans, it is unknown whether inheritance of CpG methylation patterns occurs across the genome (as in plants) or as rare exceptions (as in mammals) ^4^. Here, we demonstrate genome-wide intergenerational transmission of CpG methylation patterns from parents to sperm and larvae in a reef-building coral. We also show variation in hypermethylated genes in corals from distinct environments, indicative of responses to variations in temperature and salinity. These findings support a role of DNA methylation in the transgenerational inheritance of traits in corals, which may extend to enhancing their capacity to adapt to climate change.

---

Corals provide habitat for thousands of marine species, protect shorelines and support human livelihoods ^5^. However, as the oceans warm, repeated thermal stress events have driven corals into severe global decline ^6,7^ and the rapid rate of climate change threatens to overwhelm the capacity for adaptation by genetic means alone ^8,9^. Recent research has demonstrated that stress-induced changes in the DNA methylome of corals correlates with phenotypic changes that explain increased organismal fitness ^10,11^. Mechanistically, these changes appear to regulate transcriptional homeostasis through the inhibition of spurious transcription and transcriptional noise, reflecting adjustments of expression in response to transcriptional needs under changing conditions ^10,12^. However, due to the low heritability of DNA methylation in mammals and its complete absence in popular model organisms (e.g., *Drosophila* and *Caenorhabditis elegans*), the current perception is that this mechanism is unlikely to contribute to transgenerational plasticity, and hence to the ability of corals to respond to climate change ^9^.

Here, we initiate a paradigm shift by providing evidence for intergenerational inheritance of DNA methylation patterns in reef-building corals. The brain coral *Platygyra daedalea* ^13^ has life-history traits which we expect to promote epigenetic acclimatisation within generations, and the transfer of these modifications between generations ^8^. Namely, individual colonies are typically stress-tolerant ^14^, long lived (up to ~100 years, Supplementary Table S1), and spawn large numbers of gametes annually ^15^. Samples of *P. daedalea* were obtained from two populations in the Arabian Peninsula (Fig. 1a, b). The Abu Dhabi population (24°35’56” N, 54°’25’17” E) is highly stress-tolerant, lives under extreme temperatures (winter < 19 °C and summer > 35 °C) and salinity (40-46 psu; Supplementary Table S2), and has persisted through major thermal stress events (coral bleaching) that occurred in Abu Dhabi in 1998, 2002, 2010, and 2012 ^16,17^. In contrast, the neighbouring Fujairah population (25°29’33” N, 56°21’50” E) lives under comparatively milder temperatures (22–33 °C) with near-oceanic salinity (36–39 psu) and has not experienced anomalous temperature events in recent years.

**Fig. 1:**
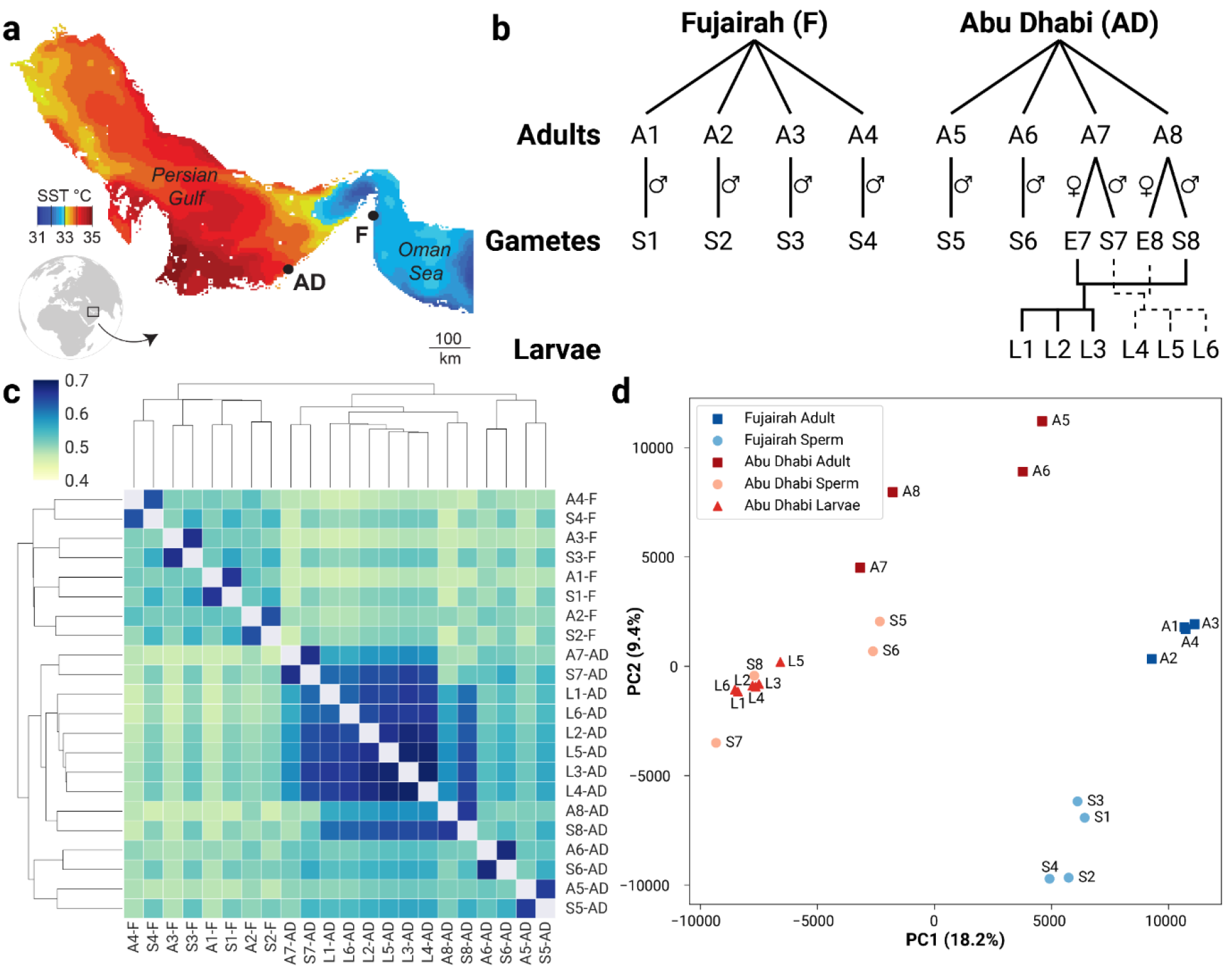
Data acquisition and summarised findings showing intergenerational inheritance of DNA methylation patterns in *P. daedalea.* (a) Contour map showing summer sea surface temperature (SST) differences between origin populations at Fujairah (F) and Abu Dhabi (AD) (mean of 5 km night-time SST for August 2015, NOAA, https://coralreefwatch.noaa.gov/satellite/bleaching5km/index.php). (b) Samples were collected from eight adult colonies (A1–A8) from Fujairah (A1–A4) and Abu Dhabi (A5–A8). Sperm (S1–S8) and eggs (E7, E8) were collected from the correspondingly numbered adult. Larval samples were produced from reciprocal crosses: three from E7 and S8 (L1–L3), and another three from S7 and E8 (L4–L6). Data from egg samples (E7 and E8) were excluded from downstream analyses due to poor qualities and insufficient coverages. (c) Clustering was performed on the pairwise correlation of methylation data from all samples. The strong effect of genotype on methylation patterns is evident from the clustering of sperm samples with their respective adults, and the larval samples with samples from their parents (A1, A8, S7, S8). The larval samples clustered equally well to siblings regardless of the direction in which crosses were performed. Additionally, samples clustered well by geographical origin (“-F”: Fujairah, “-AD”: Abu Dhabi). Colour bar shows Kendall rank correlation coefficient (τ). (d) A PCA based on the same methylated positions exhibits clear separation of samples based on geographical origin (blue: Fujairah, red: Abu Dhabi) along PC1, and by developmental stage (squares: adults, circles: sperm, triangles: larva) along PC2.

Using whole genome bisulphite sequencing (WGBS) data from 30 Illumina HiSeq2000 lanes, we identified 1.73 million CpG positions that were consistently methylated in the ~800 Mb *P. daedalea* genome (3.91% of all CpGs). These positions had a per-sample mean coverage of 45.44× across 22 samples (Supplementary Data S1b). Similar to other studied cnidarians, methylation in *P. daedalea* was predominantly in gene bodies ^10–12,18,19^. A larger fraction of CpGs in genic regions were methylated than in intergenic regions (5.0% vs. 3.6% respectively; Supplementary Fig. S1a). On a per-position basis, methylated positions in genic regions were also, on average, more strongly methylated than in intergenic regions (mean of 34.2% vs. 23.2% respectively; Supplementary Fig. S1b). Within genic regions, methylated positions were more commonly found at both ends (Supplementary Fig. S1c).

Parental genotype proved to be a strong predictor of inherited *P. daedalea* DNA methylation patterns: sperm samples were significantly more similar to their respective parents than non-parental sperm (mean τ ± SEM: 65.7 ± 0.2% vs. 53.2 ± 0.1%, *t*-test *p* < 10^−16^; Fig. 1c). This signature of inheritance was also present in the larval samples—as a group, they were significantly more similar to their parental sperm samples (S7 and S8) than to other sperm samples (S1-6; 60.5 ± 0.1% vs. 51.0% ± 0.1%, *t*-test *p* < 10^−24^). Within the group, however, the larval samples did not cluster in a pattern that reflected the reciprocal crossings, suggesting the crosses exhibit equal paternal and maternal effects. Remarkably, the variation of the methylation data captured by the first two principal components in a PCA (principal component analysis) shows a clear separation of the samples by geographical origin along PC1 (Fujairah/Abu Dhabi), and by developmental stage along PC2 (adult/sperm/larval; Fig. 1d). While the effect of genotype is not apparent in the first two principal components, PCAs of samples plotted per-location (i.e. excluding effects of geographical origin) show clustering of samples by genotype (Supplementary Fig. S2). In all PCAs, larval samples clustered closest to their parental sperm samples, and in a gender-neutral fashion.

As we were not dealing with genotypically identical samples, we wondered—to what extent—the observed epigenotypes of *P. daedalea* were determined by their individual genotypes. If epigenotypes were perfectly correlated to genotypes, it implies that the environment could only influence epigenotype via natural selection of better adapted genotypes. To estimate variation in the genotype of our samples, we analysed the WGBS reads and identified 407k SNPs present in the A/T fraction of the genome (as A and T bases remain unchanged post-bisulphite treatment). Pairwise correlations of the genetic variation were as expected: sperm samples were most similar to their respective parents, and there were fewer genetic differences between samples from the same location (Fig. 2a). If genotypes fully dictated epigenotypes, it follows that there would be a perfectly linear relationship between pairwise genotypic correlations and epigenotypic correlations. Overall, there is a strong but imperfect linear relationship (*r*^2^ = 0.805, *p* < 10^−43^) between these variables (Fig. 2b). Pairwise comparisons with the highest genetic correlations (i.e. adult vs. sperm of the same genotype) had the highest epigenetic correlation. When comparing samples across locations (Fujairah vs. Abu Dhabi, purple), they were genetically—and epigenetically—less similar than samples from the same location (blue and red), illustrating the combined effect of genotype and environment on epigenotypes. Focusing on the vertical spread of epigenetic correlation values, comparisons of adult vs. sperm samples (intermediate hues) result in correlations with lower values than adult vs. adult (dark hues) or sperm vs. sperm (light hues). This indicates that development has an effect on epigenotype.

**Fig. 2:**
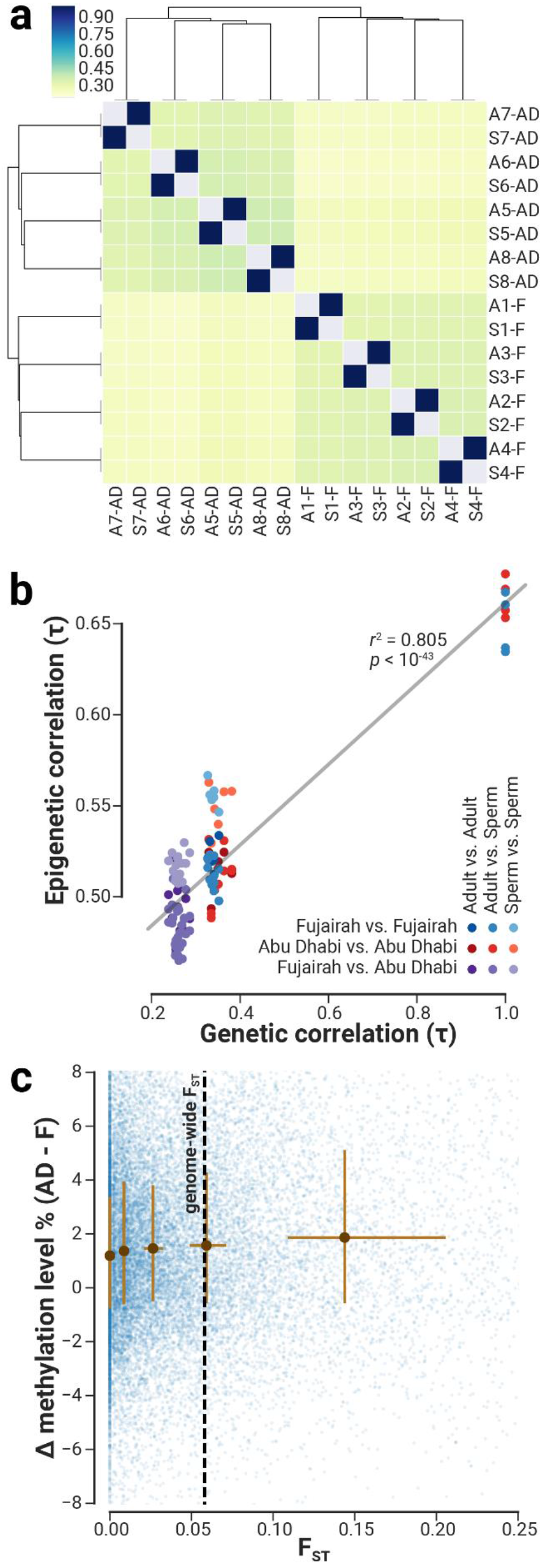
Epigenotype is strongly influenced by genotype, and to a lesser extent, by environment and development. (a) Clustering was performed on the pairwise correlation of SNP data from adult and sperm samples (*n* = 16). Samples cluster best by individual, followed by the geographical origin of the samples. Colour bar indicates Kendall rank correlation coefficient (τ values are in Supplementary Data S2a). (b) For every pairwise comparison (i.e. 16 samples: ^16^C_2_ possible pairings = 120 points), the pairwise genetic correlation (τ) is plotted on the x-axis (Supplementary Data S2b), while the pairwise epigenetic correlation (τ) is on the y-axis (Supplementary Data S2c). Points were coloured according to sample identities in the pairwise comparison (see bottom-right colour matrix). Colours are assigned based on location (blue if both are from Fujairah, red if both Abu Dhabi, purple otherwise); hues are assigned based on developmental stage (darkest hue if both are adults, lightest if both sperm, intermediate otherwise). Pairwise genetic similarities are strongly, but not completely, predictive of epigenetic similarities. The combined effects of genotype and environment contribute to the horizontal spread of points; developmental stage contributes to the vertical spread of points. (c) F_ST_S and differences in median methylation levels (Abu Dhabi - Fujairah) between populations were calculated for all genomic loci (50 kb window, 10 kb step size) in the *P. daedalea* genome. Window-specific delta methylations were plotted for windows with ≥ 60 methylated positions and ≥ 20 SNPs. As F_ST_ values were not normally distributed, they were partitioned into quintiles. Brown dots indicate median F_STS_ and median methylation differences for every quintile; brown lines indicate interquartile ranges.

To further deconvolute the separate effects of genotype and the environment on the epigenotype, we calculated F_ST_ values of genomic loci in 50 kb windows to identify regions that are genetically homogeneous/heterogeneous between the populations (Fujairah vs. Abu Dhabi). If genotype fully influences epigenotype, in genetically homogeneous loci, we expect to see approximately equal amounts of methylation in both populations (i.e. delta methylation = 0 when F_ST_ = 0); in genetically heterogeneous loci, we expect to see large differences in methylation levels between populations. Surprisingly, homogeneous loci have a median delta methylation of 1.2% (IQR: −0.7-3.4%), which significantly departs from expectations (one sample *t*-test *p* < 10^−132^). As expected, we also observe progressively larger differences in methylation levels in more heterogeneous loci (Fig. 2c). This finding demonstrates that a strong environmental effect contributes to the epigenotypes of *P. daedalea*.

We proceeded to investigate whether developmental and environmental effects had a combined effect on the epigenotypes of *P. daedalea*. Using a generalised linear model (GLM), we confirmed that genic methylation levels (*n* = 11,598 genes with ≥ 5 methylated positions) were almost always (*n* = 11,594) independently affected by development and environment, supporting the independent analysis of these effects.

We were interested in the functional significance of genes that undergo large changes in methylation levels between developmental stages. We observed generally higher methylation levels in sperm samples than their parents (median: +3.2%, mean: +5.2%; Fig. 3a), and these increases are mainly located in genic regions (Fig. 3b). As invertebrate DNA methylation is concentrated in gene bodies ^10,12,20–22^, the targeted increases in methylation levels are likely to have functional significance. To identify genes that are differentially methylated in a development-specific manner, we carried out a one-way ANOVA and filtered by effect size (corrected *p* < 0.05 and methylation levels differ > 20% between any pair of developmental stages) to produce a list of 794 genes (6.8% of all methylated genes).

**Fig. 3:**
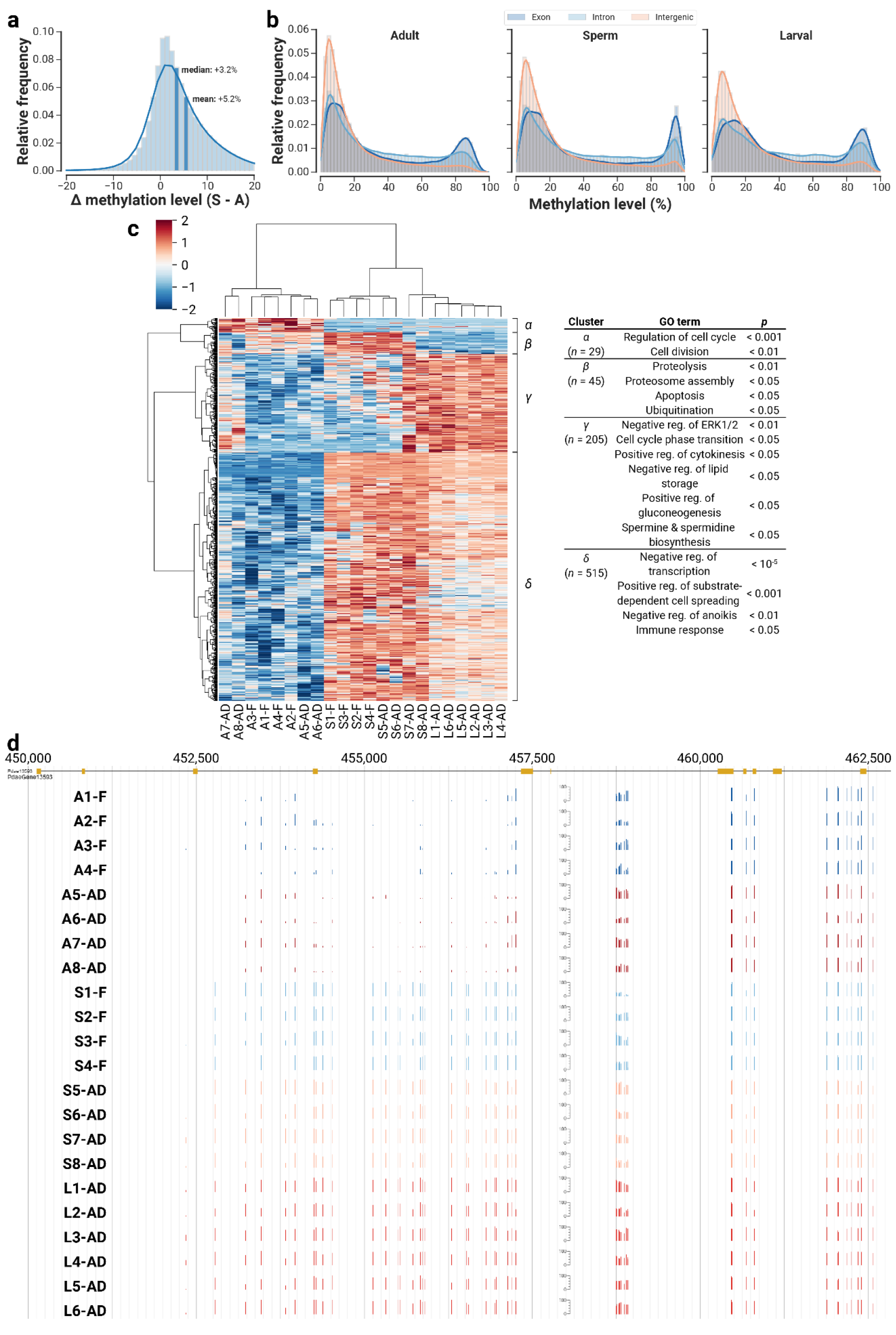
DNA methylation profiles vary considerably across developmental stages. (a) There is a general increase in per-position methylation levels in the sperm samples, relative to their parents. (b) This increase is more pronounced in genic regions (i.e. exon and introns) than intergenic regions. (c) Clustering of differentially methylated genes recapitulates the expected pattern: samples cluster best by development stage, followed by location, and larval samples are closest to their parents. Clustered genes were partitioned into four distinct groups (labelled as *α-δ*) for functional enrichment analysis: *α*, *β* and *γ* represent genes that are hypermethylated in adults, sperm and larvae respectively; while *δ* represent genes that are hypermethylated in both sperm and larvae. Values of colour bar (−2 to 2) represent z-scores. (d) An example gene (Pdae13593, putative negative regulator of anoikis) that was consistently hypermethylated among the sperm and larval samples (S1-8, L1-8).

To investigate whether genes with similar biological functions share similar methylation patterns across developmental stages, we performed a clustering analysis on the differentially methylated genes. We observed four distinct clusters: genes that are hypermethylated for a particular developmental stage and hypomethylated otherwise (clusters *α*, *β*, *γ*; Fig. 3c), and one that is hypermethylated in sperm and larvae but hypomethylated in adults (cluster *δ*, Fig. 3c). Interestingly, methylation patterns in sperm samples tend to be the intermediate of adults or larvae, suggesting that the altering of methylation levels during development is a gradual process. Regardless, our data indicates that changes in methylation could occur in the timescale of days (sperm and larval samples are 54 hours apart), in contrast to previously published studies which measured changes in samples conditioned over weeks to years (6 weeks for Putnam, et al. ^23^; 3 months for Dixon, et al. ^11^; 2 years for Liew, et al. ^10^).

Functional enrichment on individual clusters resulted in terms associated with the general biological requirements of each stage. Hypermethylated genes in adults (cluster *α*, Fig. 3c) were mostly predicted to regulate cell cycle and cell division, indicative of the different rates of cell division in adults than gametes or larval samples. The preponderance of cell lysis and ubiquitination terms in hypermethylated genes in sperm (cluster *β*, Fig. 3c) is a likely hallmark of spermatogenesis—corroborating the expanded role of ubiquitination during spermatogenesis by regulating biogenesis and stability of membranous organelles ^24^ and preventing inheritance of male mitochondria ^25^. For hypermethylated genes in larvae (cluster *γ*, Fig. 3c), the terms indicate a pro-growth phenotype. The breakdown of fatty acids, gluconeogenesis, and the synthesis of spermine and spermidine suggest increased metabolic activity, while the regulation of cell cycle and ERK proteins, a family of kinases involved in cell proliferation ^26^ suggest increased growth rates. Lastly, genes highly methylated in sperm and larvae (cluster *δ*, Fig. 3c) appear to be linked to substrate recognition (Fig. 3d shows an example gene) and symbiont recognition, both important processes that underpin successful larval settlement and symbiosis establishment ^27,28^.

In light of recent evidence for methylation changes conferring stress acclimatisation in corals ^10,11^, we were particularly interested in exploring the effect of environment on methylation levels. A similar approach with similar filters (corrected *t*-test *p* < 0.05 and pairwise methylation levels differ > 10%) identified 329 genes that were differentially methylated between Fujairah and Abu Dhabi. The choice of a lower effect size filter stems from the observation that methylation levels of samples from Abu Dhabi were not much higher than from Fujairah (median: +0.8%, mean: +1.7%; Fig. 4a). This increase was also not specific to genic regions, unlike the hypermethylation observed in sperm relative to adults (Fig. 4b).

**Fig. 4:**
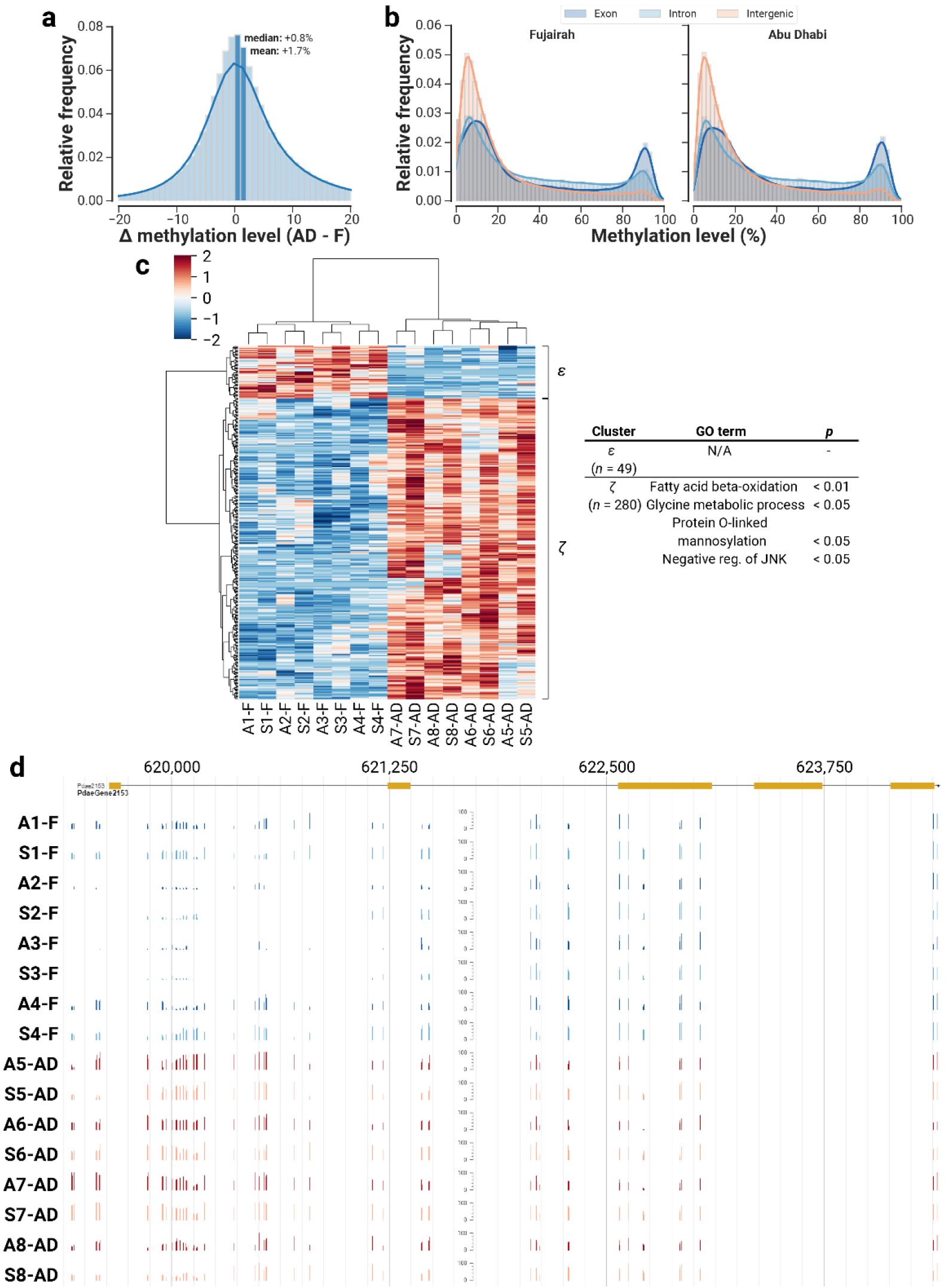
DNA methylation profiles are also associated with the environmental origin of the samples. (a) Samples from Abu Dhabi have slightly higher per-position methylation levels than those from Fujairah. (b) This increase is not confined to a specific genomic context. (c) There were fewer genes consistently hypermethylated among the Fujairah samples (cluster *ε*) than in the Abu Dhabi samples (cluster *ζ*). Notably, sperm samples clustered best with their corresponding parent, further corroborating the intergenerational inheritance of methylation patterns. Values of colour bar (−2 to 2) represent z-scores. (d) An example gene (Pdae2153, a putative negative regulator of JNK) that was consistently hypermethylated in samples from Abu Dhabi (AD). Methylation in the first intron of the gene also appears influenced by genotype.

Hypermethylated genes in Abu Dhabi (cluster *ζ*, Fig. 4c) were enriched for genes that appear to be responses to the hotter and more saline conditions of Abu Dhabi. Increased metabolism of fatty acids under heat stress is common in corals ^29,30^ and sea anemones ^31^. Glycine, in the form of mycosporine-glycine, is one of the primary mycosporine-like amino acids that confer photoprotective and antioxidative function in corals ^32^. In yeast, O-linked mannosylation has been shown to mark misfolded proteins that arise under cell stress, targeting them for degradation in the endoplasmic reticulum ^33^. The hypermethylation of JNK (c-Jun N-terminal kinase) regulators (Fig. 4d shows an example gene) echoes similar observations in the coral *Stylophora pistillata*: samples under long-term pH stress exhibited hypermethylation in the JNK pathway that were strongly predictive of the resulting phenotype ^10^. For the hypermethylated genes in Fujairah (cluster *ε*, Fig. 4c), we were unable to identify putative functions as many of those genes had no functional annotations.

In conclusion, we demonstrate—for the first time—that epigenetic modifications in reef-building corals are readily transmitted from parents to their offspring. This mode of epigenetic inheritance is similar to that of plants, instead of other well-studied metazoans (Supplementary Fig. S3). There are two possible explanations, keeping in mind conflicting molecular evidence for the existence of segregated germlines in corals ^34–36^. Firstly, the absence of a segregated germline in corals would allow for the passing down of somatic cell methylomes, akin to epigenetic inheritance in plants ^2^. For example, in *Arabidopsis thaliana*, transfer of somatic cell methylomes facilitates the retention of stress tolerance in offspring following parental stress acclimation ^37^. Alternatively, coral methylomes could escape reprogramming in the presence of a segregated germline in a similar fashion to zebrafish paternal methylomes ^38,39^. Transfer of paternal methylomes plays an important role in zebrafish embryogenesis, for example in guiding left-right asymmetry of organ positioning ^40^. Importantly, in corals, inheritance of functional DNA methylation could enable acclimatory gains in stress tolerance accumulated during long colony lifespans to be transferred to large numbers of gametes or larvae (i.e. transgenerational plasticity). These “epialleles” ^41^ could thus serve as a non-genetic substrate for Darwinian selection to act upon, and potentially accelerate evolution ^42^. By extension, long-term cultivation of corals under elevated temperatures may be a viable approach for producing fitter epialleles that could be later crossed into wild populations to enhance their resilience to rapid climate warming ^43^. Critical to these optimistic predictions is verifying that inherited epialleles do confer functional advantages to corals across generations, which could be the focus of future investigations.

## Methods

### Sample collection

*P. daedalea* colony fragments were sourced from reef sites in the Sea of Oman in the Indian Ocean (Fujairah: 25°29’33” N, 56°21’50” E; ~3 m) and the southern Persian Gulf (Abu Dhabi: 24°35’56” N, 54°25’17” E; ~6 m) which have distinct thermal and salinity profiles (Supplementary Table S2). Fragments were collected 4-6 nights prior to the full moon on 4 May 2015 and housed in 1,000-litre recirculating aquarium systems at New York University Abu Dhabi at the ambient temperature and salinity measured at the time of collection (F: 27.6 °C and 35.7‰; AD: 27.9 °C and 41.2‰). *P. daedalea* is a simultaneous hermaphrodite and broadcast spawning activity was observed from −4 nights to +10 nights relative to the full moon, with most individual fragments releasing gametes over several nights ^44^. During this period, samples were collected from adults (F: 4; AD: 4), sperm (F: 4; AD: 4), eggs (AD: 2), and larval crosses (AD: 2), snap-frozen and stored at −80 °C. Prior to spawning, individual coral fragments were isolated in buckets to prevent cross-contamination and fertilization. Gamete bundles (eggs + sperm) were collected with transfer pipettes, sperm was isolated by filtration (60 μm plankton mesh) and concentrated by centrifugation. Larvae were bred from reciprocal crosses of gametes from two Abu Dhabi colonies, reared in 1 μm filtered seawater in triplicate cultures, and were sampled at 54 hours post-fertilization (i.e., planula stage of development). This design enabled us to evaluate both population (Fujairah vs. Abu Dhabi) and intergenerational (adults vs. sperm vs. larval) variation in DNA methylation profiles.

### Preparation and sequencing of whole genome bisulphite libraries

DNA were extracted from all samples by suspending them in 750 μl of buffer (100 mM Tris/EDTA/NaCl, 1% SDS) preheated to 65 °C. After 3-4 hours incubation, RNA contaminants were removed by addition of 1 μl of RNase A (10 μg, 15 min). Extracts were chilled, proteins were removed (1 M potassium acetate), DNA was precipitated (100% isopropanol), washed (70% ethanol), re-suspended (0.1 M Tris; 200 μl for adults, 100 μl for sperm and 20 μl for eggs), and stored at 4 °C.

Bisulphite DNA libraries were prepared using NEBNext Ultra II DNA Library Prep Kit for Illumina (NEB, Ipswich, MA), following manufacturer’s instructions. Methylated TruSeq Illumina adapters (Illumina, San Diego, CA) were used for adapter ligation. Bisulphite conversion with the EpiTect Bisulfite Kit (Qiagen, Hilden, Germany) was carried out with cycling conditions of 95 °C for 5 min, 60 °C for 25 min, 95 °C for 5 min, 60 °C for 85 min, 95 °C for 5 min, 60 °C for 175 min, then 3 cycles of 95 °C for 5 min, 60 °C for 180 min, and held at 20 °C for ≤ 5 hours. Subsequently, libraries were enriched with KAPA HiFi-HotStart Uracil+ ReadyMix PCR Mix (KAPA Biosystems, Wilmington, MA), following manufacturer’s instructions. Libraries were quality checked on a Bioanalyzer DNA 1K chip (Agilent, Santa Clara, CA) and quantified using Qubit 2.0 (Thermo Fisher Scientific, Waltham, MA), prior to pooling in approximate equimolar ratios for sequencing.

Pooled libraries were initially sequenced on 24 lanes of HiSeq2000 (Illumina, San Diego, CA). Based on the results of the sequencing, libraries with lower coverages were resequenced on 6 additional lanes of HiSeq2000 (Supplementary Data S1a).

### Identification of methylated positions

From both runs (i.e., 30 lanes), a total of 4.13 billion read pairs were obtained across 24 samples, and subsequently trimmed using cutadapt v1.8 ^45^ (Supplementary Data S1a). Following trimming, reads were mapped to the draft *P. daedalea* genome (REF), deduplicated and scored on a per-position basis for methylated and unmethylated reads using Bismark v0.17 ^46^. At this stage, we excluded data from the egg samples (*n* = 2) from subsequent analyses as the data from these samples were very poor (struck-out red rows in Supplementary Data S1b).

To ensure that methylated positions were *bona fide*, four stringent filters were applied. Firstly, on the pooled dataset, the probability of methylated positions arising from chance on a per-position basis was calculated using a binomial distribution of B(*n*, *p*), where *n* represents total coverage (methylated + unmethylated reads) and *p* the probability of sequencing error (set to 0.01 to mimic a Phred score of 20). Positions with *k* methylated reads were kept if *p*(X ≥ *k*) < 0.05 (post-FDR correction). Secondly, positions were kept only if there were at least a methylated read in all biological replicates of at least one biologically meaningful group (i.e., *n* = 3 or 4 Fujairah adults, Fujairah sperm, Abu Dhabi adults, Abu Dhabi sperm or Abu Dhabi larvae). Thirdly, positions were retained only if median coverage was ≥ 10 and minimum coverage was ≥ 5 across all 22 samples. These filtering steps were very conservative: 20.6 million positions had at least 1 methylated read mapping to it, but only 1.7 million positions passed all filters. The scripts used for this section (and further theoretical justifications) are available at https://github.com/lyijin/working_with_dna_meth.

### Assignment of genomic context to methylated cytosines

A Python script was used as previously described ^10^. Briefly, the script reads the GFF3 annotations of the *P. daedalea* genome and the positional coordinates of the methylated cytosines to assign genomic context to the methylated position. Distances to the 5’ and 3’ end of each genomic feature (gene/intergenic region/exon/intron) were calculated for downstream analyses.

Overall, 16,035 genes (64.2% of all genes) had at least one assigned methylated position; 12,111 genes (48.5%) had ten or more assigned methylated positions.

### Identification of SNPs in the *P. daedalea* genome

Using the deduplicated BAM files produced from the Bismark pipeline (all adult and sperm samples, *n* = 16), coverages across all bases were compiled into a table via samtools (“samtools mpileup”, with the *P. daedalea* genome providing the per-position reference base). Reference bases corresponding to C and G were discarded, as bisulphite treatment converts C-to-T (and G-to-A on the opposite strand).

A Python script further imposed three rules to filter for potential SNPs that are highly covered: across all 16 samples, median coverage at that position must be ≥ 20, and the minimum coverage ≥ 10. The overall minor allele frequency (MAF), calculated by dividing the sum of all non-genome-matching bases over total coverage (i.e. values range from 0 to 1 inclusive), must be between 5% and 95%. A value below 5% implies that the non-genome-matching bases probably arose from sequencing errors; a value above 95% is likely due to the reference base being wrongly called. If a position passes these three criteria, MAFs were calculated on a per-sample basis and stored as an intermediate file.

Using another Python script, per-position genotypes were assigned based on the MAFs, as listed below:

MAF ≤ 25%: genotype 0 (i.e. homozygous reference base)
25% < MAF ≤ 75%: genotype 1 (i.e. heterozygous)
MAF > 75%: genotype 2 (i.e. homozygous non-reference base)

As we assumed that the adult and sperm samples from the same individual should be genetically identical, positions were only retained if all sperm samples and their respective parents had the same genotype (i.e., A1–8, when written out as a list, would be identical to S1–8 written out in a similar fashion). A total of 407,419 positions passed this final filter, and were deemed SNPs.

### Calculation of Kendall ranked correlation coefficients for pairwise comparisons

As both methylation data and SNP data were not normally distributed, we opted to calculate the Kendall ranked correlation coefficients (τ) to measure whether data were similarly ordered in a similar fashion in datasets *X*and *Y*. The range of τ spans [−1, 1], where −1 indicates perfectly discordant ranked observations; 1 indicates perfectly concordant ranked observations.

Correlation coefficients were calculated using Python scripts, which enabled the plotting of Figs. 1c, 2a and 2b. For transparency, these values are tabulated in Supplementary Data S2.

### Assessing effect of genotype on epigenotype

To assess the effect of genetic heterogeneity on DNA methylation levels, the *P. daedalea* genome was partitioned into 50 kb windows with a step size of 10 kb. Genetic heterogeneity was estimated using F_ST_—but as there are various ways to calculate this value, we adopted Hudson’s definition and calculated per-position F_ST_ values using equation 10 described in Bhatia, et al. ^47^. The paper also recommended that F_ST_S should be averaged in a window as a ratio of averages, using the equation (as derived in the Supplementary Materials of Bhatia, et al. ^47^ but reproduced here for convenience):

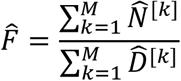

where *F* is the averaged F_ST_ value of a particular 50 kb window and markers *k* = 1, 2, …, M denote individual SNPs within this window. *N* and *D* are the numerator and denominator of Hudson’s equation respectively:

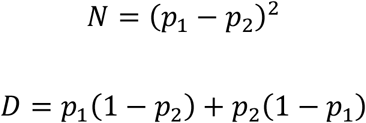

where *p*_1_, *p*_2_ denote allelic frequencies of the non-reference base among the Fujairah and Abu Dhabi populations respectively.

The epigenotypic variances within the same 50 kb windows were calculated by first calculating the window-specific mean methylation levels across all Fujairah samples, and again for all Abu Dhabi samples. The absolute difference between these two means thus indicates the extent of the differences in methylation levels between the two populations.

As the draft genome still contains regions of unknown bases (Ns), we retained windows that had ≥ 20 SNPs and ≥ 60 methylated sites. Correlations of these computed values were all implemented in Python.

### Identification of differentially methylated genes

Individual positions have methylation levels that range from 0 (not methylated at all in all cells) to 1 (completely methylated in all cells). To extend this measure to genes, which contain multiple methylated positions, medians of the methylation levels were calculated for every gene (medians are less prone to outliers than means). Furthermore, to reduce noise, we focused on genes that had at least five or more methylated positions, filtering out 3,068 genes that did not meet this criterion. The genes that were filtered out were mostly unannotated, or annotated as proteins with unknown functions.

To ensure appropriate statistical models were applied for the analysis of our dataset, we first checked whether genic methylation levels were normally distributed within biologically meaningful groups ^48^, and whether variances were equal between groups ^49^. For the vast majority of genes, the null assumptions of normality and equal variance could not be rejected (Supplementary Data S3). Hence, we used untransformed methylation levels in all subsequent statistical tests.

Genic methylation levels were thought to be influenced by two variables: developmental stage (“devt”: adult/sperm/larvae) and sample origin (“origin”: Fujairah/Abu Dhabi). As the interaction of both variables could confound statistical analysis that focused on individual variables, a GLM (generalised linear model) was implemented in Python with the general equation:

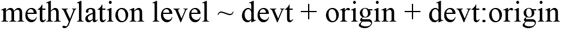

Our results indicate that methylation levels of only 4 of 11,598 genes had a significant interaction between developmental stage and origin (Supplementary Data S4). Consequently, we infer that methylation level are independently affected by developmental stage and sample origin, justifying the separate analysis outlined in Supplementary Fig. 4.

To identify genes that were differentially methylated due to developmental stage, one-way ANOVAs were carried out on a per-gene basis encompassing three developmental stages (*n* = 8 adults, 8 sperm, 6 larvae). As many genes were considered significant (3,863 of 11,598), we filtered for genes with a methylation level change of > 20% in any direction between any pair of developmental stages. This conservative filter picked out 794 genes that exhibited significant and large changes during development (Supplementary Data S5).

To identify genes that showed significant differences in methylation level due to geographical origin, *t*-tests were carried out on a per-gene basis for all Fujairah samples against non-larval Abu Dhabi samples (*n* = 8 Fujairah, 8 Abu Dhabi). Comparatively fewer genes (406 of 12,111) were considered significant. As samples from the different origins had similar methylation levels, a less stringent effect size filter of > 10% was applied, resulting in a final set of 329 genes that exhibited significant and large changes between locations (Supplementary Data S6).

Clustered heatmaps were plotted for these two classes of differentially methylated genes with seaborn (https://github.com/mwaskom/seaborn), a plotting library in Python. Non-default parameters include the choice of clustering method (Ward, instead of UPGMA), and the use of z-score values (i.e. individual values reflect the number of standard deviations away from the mean).

### Multiple testing correction of *p* values

Unless otherwise stated, computed *p* values were subjected to the Benjamini-Hochberg-Yekutieli multiple testing correction ^50,51^.

### Functional enrichments within genes of interest

GO term annotations were obtained from the draft *P. daedalea* genome (REF). topGO ^52^ was used, with default settings, to perform GO term enrichment analyses on the genes of interest list. Multiple testing correction was not applied on the resulting *p*-values as the tests were considered to be non-independent ^52^. GO terms with *p* < 0.05 and occurring ≥ 5 times in the background set were considered significant (Supplementary Data S7).

## Raw data

Whole genome bisulphite sequencing data can be found in NCBI BioProject PRJNA430328. Individual SRA accessions are listed in Supplementary Data S1c.

(Note: data is currently under embargo, and will appear as if it hasn’t been uploaded on the NCBI website. This embargo will be lifted upon publication)

## Code availability

Python scripts and key input files are available at https://github.com/lyijin/pdae_dna_meth. Explanatory notes for the scripts can be found in the repository.

## Acknowledgements

We thank D. Abrego, G. Vaughan, and D. McParland for assistance with fieldwork, coral spawning, and the collection of environmental data, as well as the KAUST Sequencing Core Facility for the sequencing of the libraries. This work was financially supported by KAUST (sequencing and bioinformatics) and NYUAD (fieldwork and coral spawning).

## Contributions

E.J.H, Y.I. and M.A. conceived and coordinated the project. M.A., X.W. and J.A.B provided resources. E.J.H. collected samples from the wild, performed controlled crosses, and extracted DNA from fixed samples. C.T.M. constructed libraries for WGBS and RNA-seq. Y.J.L., Y.I. and M.A. analysed data. Y.J.L., E.J.H. and M.A. wrote the manuscript. All authors read and approved the manuscript.

## Supplementary information

**Supplementary Fig. S1:**
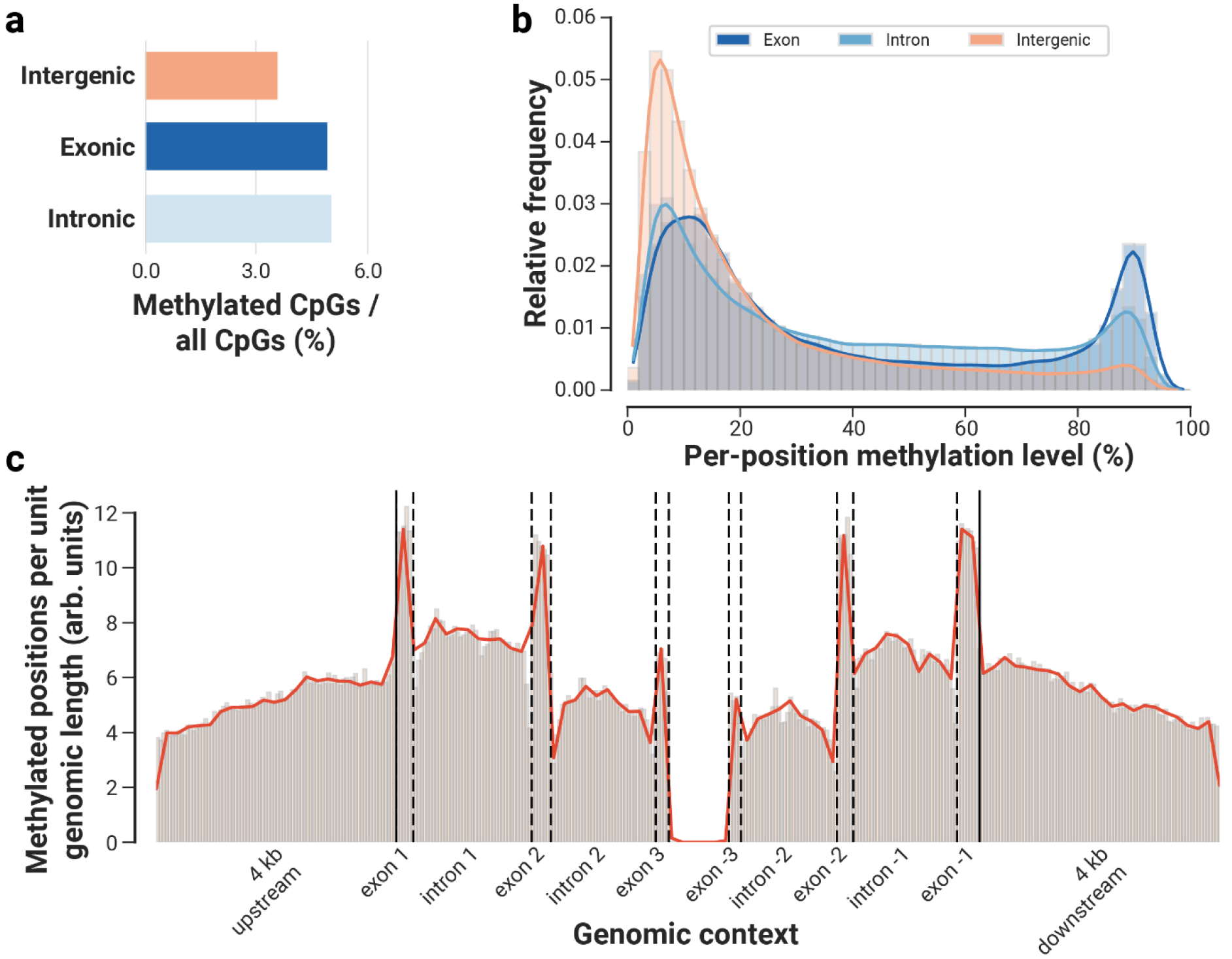
Methylation in *P. daedalea* is more commonly found in genic regions, and concentrated closer to the 5’ and 3’ ends. (a) Genic regions are more frequently methylated (~5.0% CpGs are methylated) than intergenic region (3.6%). (b) Methylation levels are bimodally distributed in exons, introns and intergenic regions. Exons have the highest methylation levels, followed by introns and intergenic regions. (c) Relative frequencies of methylated positions across a standardised gene model with flanking 4 kb regions indicate that methylated positions are more frequently found at both ends of the gene model. Solid lines depict transcriptional start site (left) and transcription termination site (right) while dotted lines delineate the borders of the indicated genomic feature. The widths of the features correspond to mean normalised lengths of the respective exons and introns in *P. daedalea* (exons, from left to right: 286 bp, 320 bp, 225 bp, 203 bp, 270 bp, 380 bp; introns, from left to right: 1,971 bp, 1,744 bp, 1,598 bp, 1,728 bp).

**Supplementary Fig. S2:**
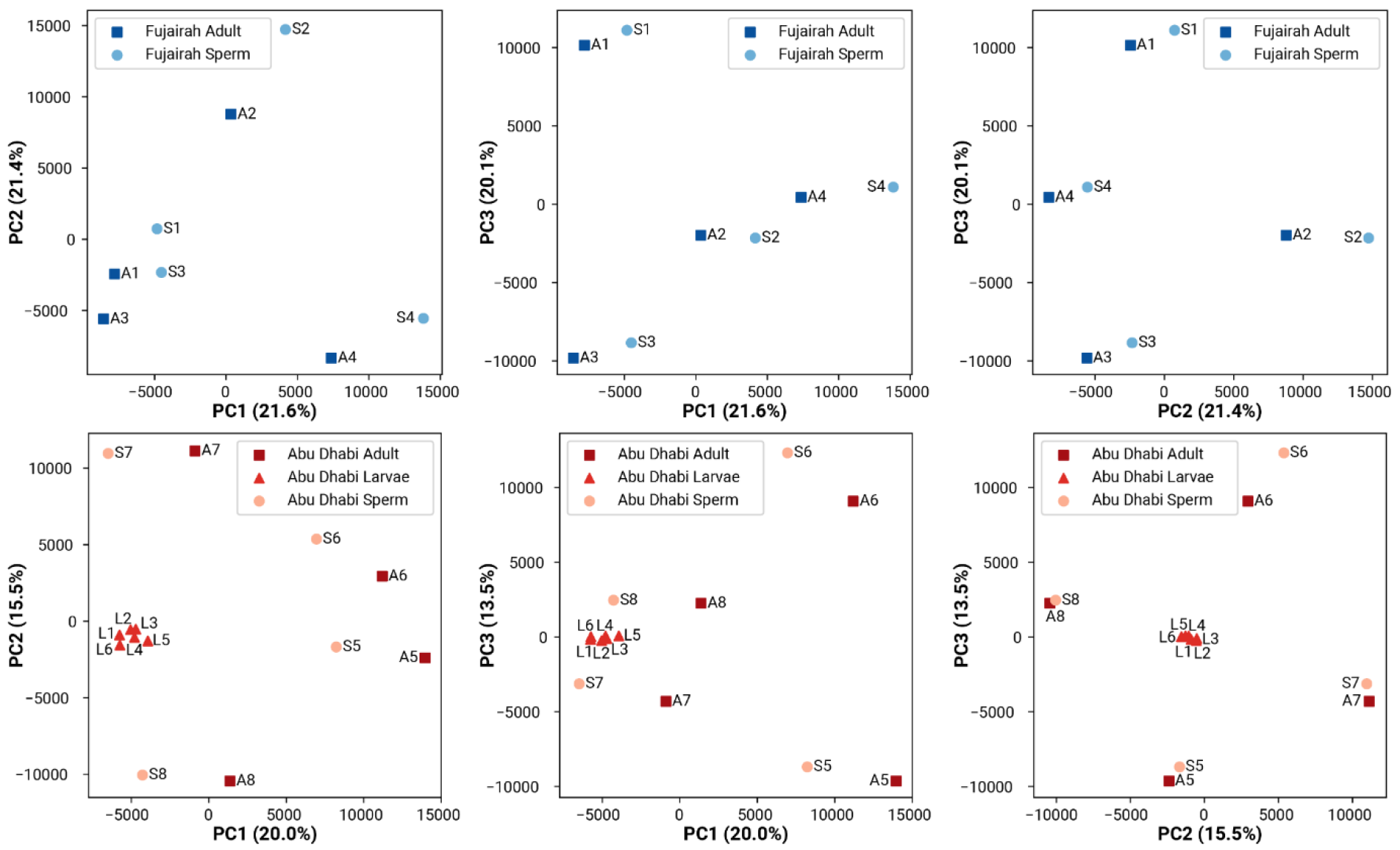
Per-location PCAs of DNA methylation patterns in *P. daedalea*. PCAs were carried out on Fujairah (top row) and Abu Dhabi samples (bottom row) separately on the first three principal axes. Samples tended to pair by genotype. The larval samples (red triangles) are positioned approximately midway of sperm samples S7 and S8 along all plotted axes, suggesting equal contribution from both parents to their DNA methylation patterns.

**Supplementary Fig. S3:**
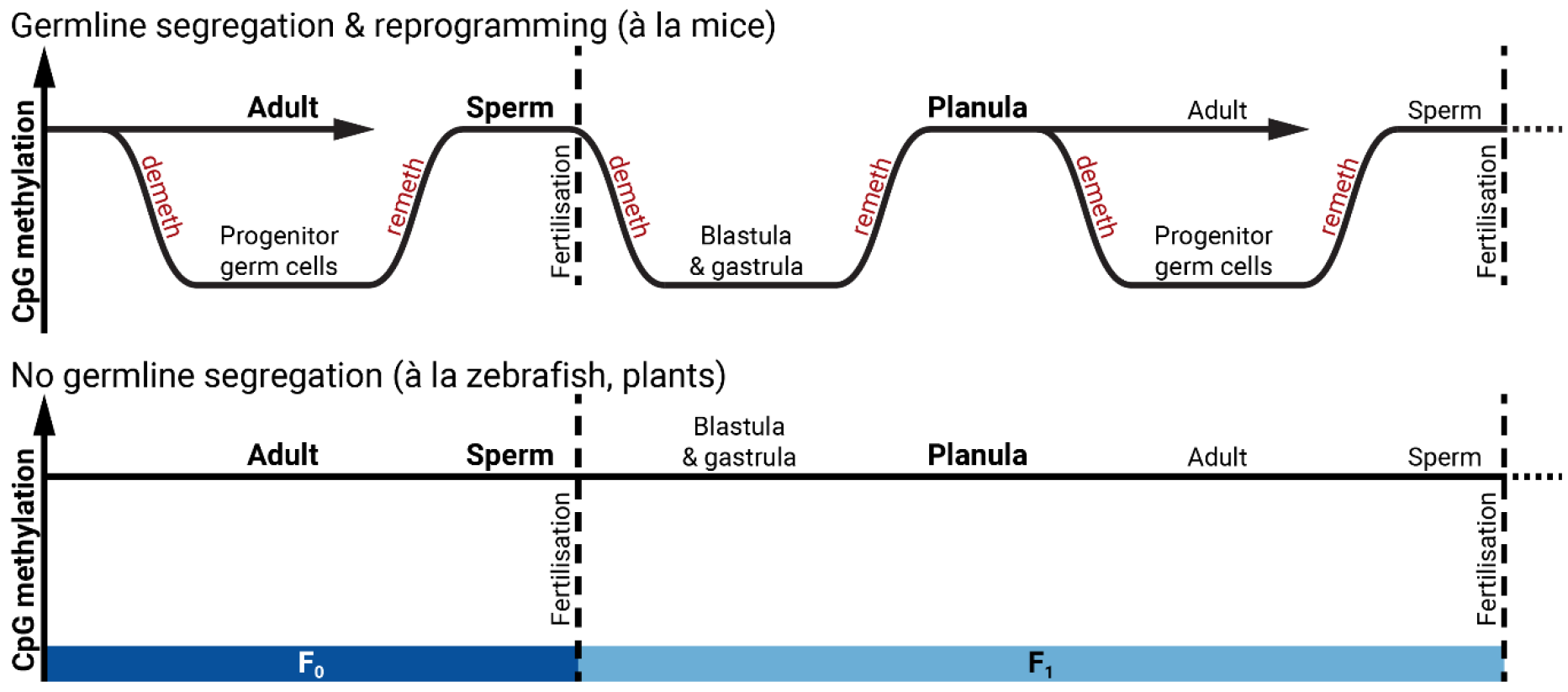
Transgenerational inheritance of CpG methylation in corals modelled after mammalian or plants systems. Bolded text (“adult”, “sperm” and “planula”) indicates equivalent samples for which we have methylation data in reef-building corals. Consistent with both models, methylation levels from all of our samples are relatively high. However, epigenetic reprogramming (twice per generation) in mice prevents the transgenerational inheritance of most epigenetic variation acquired in adults; on the other hand, CpG methylation are not reprogrammed in plants.

**Supplementary Fig. 4:**
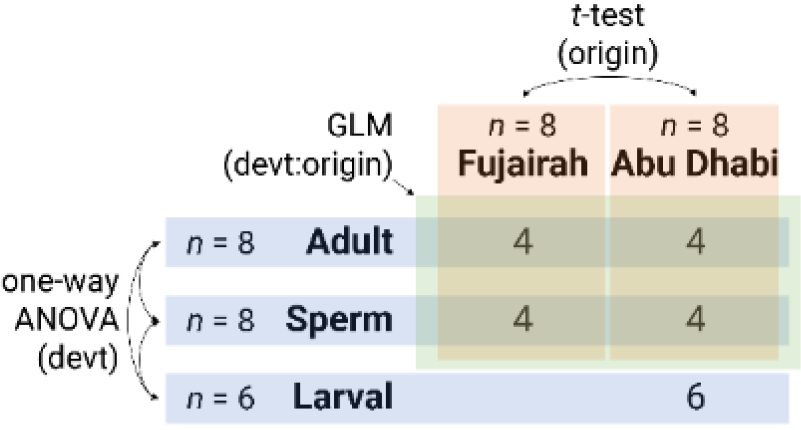
Outline of statistical tests performed. Numbers denote biological replicates of different developmental stages and sample origins, while boxes denote the groupings used in the statistical tests. The initial GLM (green) tested whether both variables had significant interaction. Subsequently, the one-way ANOVA (blue) identified genes that were differentially methylated across developmental stages, while the *t*-test (red) identified genes that were differentially methylated across sample origins.

**Supplementary Table S1.** Age estimates of *Platygyra daedalea* colonies from Abu Dhabi. Colony radius-age relationships established for *P. sinensis* from the Great Barrier Reef ^53^ were applied to local data as this closely related species has identical growth morphology to *P. daedalea*. Radial size gains were adjusted to reflect a 33% slower growth rate in Abu Dhabi *P. daedalea* ^54^ and applied to the sizes of colonies measured in the population (*n* = 140).

|  | *P. sinensis*<br>growth rates<br>(years per cm) | *P. daedalea*<br>growth rates<br>(years per cm,<br>adjusted) | *P. daedalea*<br>colony radius<br>(cm) | *P. daedalea* colony<br>age estimates<br>(mean, min–max) |
| --- | --- | --- | --- | --- |
| **Mean** | 1.9 | 2.5 | 12.8 | 32 (24–41) |
| **Minimum** | 1.4 | 1.9 | 1.25 | 3 (2–4) |
| **Maximum** | 2.4 | 3.2 | 47.8 | 120 (91–153) |

**Supplementary Table S2.** Temperature and salinity profiles of *P. daedalea* populations in Abu Dhabi and Fujairah. Temperatures summaries were calculated from daily means from *in situ* temperature loggers (*HOBO* pendant) attached to the reef substrate (Abu Dhabi: 2010-2014; Fujairah; 2012-2014). Salinity summaries were calculated from ad hoc measurements (YSI multi-probe) taken *in situ* within 0.5 m of the substrate (Abu Dhabi: 2013-2017, *n* = 23; Fujairah; 2013-2015, *n* = 9).

| **Temperature (°C)** | **Abu Dhabi** | **Fujairah** |
| --- | --- | --- |
| Annual mean | 28.1 | 27.4 |
| Maximum | 35.7 | 33.3 |
| Minimum | 18.4 | 22.0 |
| Annual range | 17.3 | 11.3 |
| **Salinity (psu)** | **Abu Dhabi** | **Fujairah** |
| Mean | 42.1 | 37.7 |
| Maximum | 45.9 | 39.4 |
| Minimum | 39.6 | 35.5 |

